# Strong breeding colony fidelity in northern gannets following High Pathogenicity Avian Influenza Virus (HPAIV) outbreak

**DOI:** 10.1101/2023.05.02.539030

**Authors:** David Grémillet, Aurore Ponchon, Pascal Provost, Amandine Gamble, Mouna Abed-Zahar, Alice Bernard, Nicolas Courbin, Grégoire Delavaud, Armel Deniau, Jérôme Fort, Keith C. Hamer, Ruth Jeavons, Jude V. Lane, Liam Langley, Jason Matthiopoulos, Timothée Poupart, Aurélien Prudor, Nia Stephens, Alice Trevail, Sarah Wanless, Stephen C. Votier, Jana W.E. Jeglinski

## Abstract

High pathogenicity avian influenza virus (HPAIV) caused the worst seabird mass-mortalities on record in Europe across 2021-2022. The northern gannet (*Morus bassanus*) was one of the most affected species, with tens of thousands of casualties in the northeast Atlantic between April-September 2022. Disease outbreaks can drastically modify the movement ecology of animals and diminish spatial consistency, thereby increasing the potential for disease transmission. To detect potential changes in movement behaviour, we GPS-tracked breeding adults following the initial HPAIV outbreak, at three of the largest gannet breeding colonies where major mortality of adults and chicks occurred (Bass Rock, Scotland, UK; Grassholm, Wales, UK; Rouzic island, Brittany, France). Crucially, GPS-tracked birds remained faithful to their breeding sites and did not prospect other breeding colonies. They performed regular foraging trips at sea, similar to their behaviour before the outbreak. Gannet foraging effort was nonetheless lower than in 2019, thus surviving birds may have benefited from reduced intra- and interspecific food competition. Breeding colony fidelity of adult northern gannets following HPAIV mass-mortalities suggests limited long-term capacity to virus spread, which may contrast with the behaviour of adults during the disease outbreak, or with that of younger individuals.

## Introduction

Avian influenza was first reported in farmed poultry in 1878 in Northern Italy (Perroncito, 1878). Outbreaks of the virus occurred worldwide in domestic fowl across the 20^th^ century (Swayne and Suarez, 2000). During this period, wild birds were also regularly affected (Gauthier-Clerc et al., 2007). In 1996, a form of highly pathogenicity avian influenza virus (HPAIV; H5Nx clade 2.3.4.4) evolved in farmed poultry in Southern China. It subsequently spilled over to wild birds, and occasionally to humans (Alexander and Brown, 2009). HPAIV has since occurred in 61 countries and is endemic in poultry farms around the world. In 2021/2022, the worst-ever HPAIV outbreak was recorded in poultry farms, especially in Europe (Wille and Barr, 2022), resulting in the culling of an estimated 47.7 million birds (Authority et al., 2022). These events deeply challenge intensive bird farming, as high-density housing conditions and long-distance poultry transport lead to the spread of HPAIV, and to the suffering and slaughter of captive animals (Kuiken and Cromie, 2022). Rapid evolution in high-density, confined poultry farms also enhances the likelihood of mutations leading to the emergence of HPAIV as the next human pandemic (Roberts Jr and Krilov, 2022). Finally, HPAIV is now causing substantial mortalities to wild birds, including in species already challenged by other human activities (Gamarra-Toledo et al., 2023).

This is the case for seabirds, which stand out as the most endangered bird group beyond parrots (Dias et al., 2019). Half of the worldwide abundance of seabirds has been lost since 1970 (Grémillet et al., 2018), mainly because of breeding habitat loss, mortality caused by fishing gear, invasive species (e.g. cats and rats), food competition with fisheries, direct and indirect consequences of climate change, and pollution (Dias et al., 2019; Grémillet et al., 2018). Pathogens can also have significant impacts on seabird populations, as demonstrated by avian cholera (*Pasteurella multocida*) affecting Cape cormorants (*Phalacrocorax capensis*) in South Arica (Crawford et al., 1992), or common eiders (*Somateria mollissima*) in North America (Iverson et al., 2016). While seabirds are common hosts of low pathogenicity avian influenza viruses (LPAIV) (Lang et al., 2016), HPAIV has rarely been detected in this group before the 2021-2022 outbreak, except in Southern Africa where recurrent HPAIV outbreaks have affected seabirds since 2016 (Molini et al., 2020). Now, for the first time, HPAIV is significantly affecting European seabird populations, with the virus spreading to e.g. 12 species in the UK (Falchieri et al., 2022). With tens of thousands of casualties in 2022 in the Northeast Atlantic, northern gannets (*Morus bassanus*) are among the most affected species, together with great skuas (*Stercorarius skua*) and sandwich terns (*Thalasseus sandvicensis*). Gannets are an iconic species that attract conservation and media interest, in their own right, but also as umbrella species for the marine environment. Despite living in some of the most anthropogenically modified marine regions of the planet (e.g. the North Sea), gannet populations consistently increased across their distribution zone in the northeast Atlantic during the second half of the 20^th^ century (Grandgeorge et al., 2008) - until the 2022 HPAIV outbreak.

The present HPAIV panzootic highlights the urgent need for a better understanding of the impact of such disease outbreaks on the movement ecology of affected species, both during and following HPAIV outbreaks, and the associated consequences for disease transmission. Seabirds such as gannets are organised as metapopulations where colonies are connected through dispersal movements of pre-breeding birds (Jeglinski et al., 2022). In contrast, breeding seabirds generally show very high fidelity to their colony and to their breeding site (Hamer et al., 2001). As central place foragers, they travel large distances away from their breeding colony to their foraging sites at sea, but many species show a marked degree of repeatability in their individual movement and space use patterns (Guiry et al., 2022). Gannets in particular have also been shown to forage in colony-specific non-overlapping areas (Wakefield et al., 2013) suggesting a limited capacity for close contact (and thus potential transmission routes) between breeders from different colonies. However, it is currently unclear how disease and the disruption caused by a disease outbreak affect the predictable movement patterns of seabirds, both in the short-term (i.e. during a disease outbreak) or in the long-term (i.e. following a disease outbreak).

Bird aggregations, especially dense seabird colonies, function as information centres (Evans et al., 2016) where birds glean public information from their conspecifics, such as the location of potential food resources at sea (Courbin et al., 2020), or the quality of local breeding habitats (Danchin et al., 1998). During the 2022 HPAIV outbreak, when many chicks and adult gannets died at breeding colonies, surviving adult gannets were exposed to public information signalling that they were breeding in a highly hazardous habitat. As a short-term effect of such disturbance, we might expect to see drastic changes in movement and space use patterns. Indeed, recent work has demonstrated that adults black-legged kittiwakes (*Rissa tridactyla*) that perceived their breeding habitat as poor immediately modified their movement patterns (Ponchon et al., 2017): Adults which failed breeding thereby switched from exclusively commuting between their nest and foraging areas at sea, to visiting other breeding colonies to prospect for better nesting spots. Prospecting behaviour was also recorded in 12 out of 14 gull and tern species studied in Europe and North America, even when individuals were successfully breeding (Kralj et al., 2023). Such changes in movement patterns increase the potential for disease transmission, by increasing the likelihood of contact, be it on land or at sea, between birds from different colonies (Boulinier, 2023; Boulinier et al., 2016).

Further, we do not know whether a disease outbreak has longer lasting effects on movement and space use of surviving animals. For example, seabird mass-mortalities are likely to reduce food competition for surviving individuals and may therefore lead to modified foraging patterns. Short-term effects on movement, such as the increase in prospecting frequency described above, could become a more permanent feature. This would lead to increased connectivity between usually segregated breeding colonies and thus have the potential to maintain enhanced disease transmission, within gannet metapopulations and at the scale of multispecies seabird communities (Boulinier et al., 2016). Finally, survivors could make permanent decisions regarding their terrestrial space-use by abandoning their breeding colony, as shown in Cape cormorants or Double-crested cormorants (*Phalacrocorax auritus*) which desert breeding colonies entirely when exposed to avian cholera or Newcastle disease virus (Crawford et al., 1992; Leighton et al., 2021). It is uncertain whether gannets display such short or long-term behavioural shifts, because they are known for their high fidelity to nesting (Nelson, 1966), and foraging sites (Votier et al., 2017; Wakefield et al., 2015). Also, studies on closely-related cape gannets (*Morus capensis*) showed that, even when confronted with strongly degraded environmental conditions and ecological traps, adult breeders kept the same breeding and foraging locations (Grémillet et al., 2016; Pichegru et al., 2010).

Previous investigations have described the movement ecology of seabirds, e.g. Indian yellow-nosed albatrosses (*Thalassarche carteri)* (Ponchon et al., 2021) or brown skuas (*Stercorarius antarcticus*)(Lamb et al., 2022) exposed to avian cholera. Yet, to the best of our knowledge, no study has investigated seabird movements following an HPAIV outbreak, and very few did so in other birds (Gaidet et al., 2010). We used the rare opportunity to fill this critical gap in three of the largest gannet breeding colonies in the Northeast Atlantic which experienced very high levels of HPAIV-related chick and adult mortality; the Bass Rock in Scotland (UK), Grassholm in Wales (UK), and Rouzic Island in Brittany (France). Bass Rock was, before the HPAIV outbreak, the largest northern gannet colony in the world, with an estimated 75,259 breeding pairs (Murray et al., 2014). Taken together, the three studied colonies totalled ca. 26% of the northern gannet world population (134,259 out of 525,694 pairs (Murray et al., 2015)).

We focussed on the longer-term effect of the HPAIV outbreak on breeding gannet space use, since potential persistent behavioural shifts may maintain a high degree of transmission potential in the metapopulation (Boulinier, 2023). We also provide some background epidemiological information and data on gannet colony dynamics during the outbreak. The outcome of our study has strong implications for understanding the potential of breeding gannets as vectors of HPAIV.

## Methods

Northern gannets were studied in May-October 2022 on Bass Rock, Scotland (56°04’N, 2°29’E), Grassholm, Wales (51°43′N, 05°28′W), and Rouzic Island, Brittany (48°54’N, 3°26’W). All personnel involved in field operations were equipped with full-body suits and disposable gloves, face masks, goggles and caps. Biocides (Virkon® or SafeFour®) were used between each manipulated bird to clean clothing, shoes and equipment.

### Characterization of the HPAIV outbreak

#### Bass Rock

On 4^th^ June 2022, unusual numbers of dead and dying gannets were observed on Bass Rock (Scottish Seabird Centre, pers. comm., see also Suppl. 1). Due to permit issues, gannets equipped with GPS-GSM tags (see below) were not screened for HPAIV.

#### Grassholm

Following an unusually high number of dead gannets at the colony, HPAIV was confirmed from Grassholm on 3^rd^ August with no further access to site permitted for the rest of the breeding season.

#### Rouzic

Mortality affecting adult gannets was first recorded on 1^st^ July 2022 on the live camera monitoring the Rouzic colony. Samples were subsequently collected from 31 live or dead gannets in, and around the Rouzic colony between July and September 2022, for HPAIV screening and characterization. We thereby performed opportunistic necropsies on four adult birds found dead on the colony, from which we collected tracheal and cloacal swabs and/or lung and trachea samples. We also collected cloacal swabs, tracheal swabs and/or external wipes impregnated with sterile distilled water (applied to the feathers and featherless areas of the birds) from live individuals showing no HPAIV symptoms, including 15 adult birds equipped with GPS-GSM loggers (see below), and 12 recently fledged juvenile birds captured at sea or in the colony. We also collected plasma samples from seven of these juvenile birds. All samples were stored at -20°C until laboratory analyses.

Samples were analysed using the protocols recommended by the French Ministry of Agriculture and based on the recommendations provided by the OIE/FAO international reference laboratory for avian influenza. Tissue and swab samples were screened for HPAIV genetic material by real-time reverse transcription polymerase chain reaction (rRT-PCR) targeting the M gene. To identify the subtype of the circulating virus, positive samples were then typed using an rRT-PCR targeting either the H5 or H7 type of the haemagglutinin gene. One sample testing positive for H5 was analysed for further characterization of the virus with an RT-PCR discriminating HP H5 of the 2.3.4.4b clade, and an RT-PCR targeting the N1 type of the neuraminidase gene. Samples testing negative for host genetic material were classified as inconclusive, likely due to the degradation of the samples, and were excluded from the study. Plasma samples were screened for antibodies specifically targeting H5N1, H5N3, H5N5 or H5N8 by haemagglutination inhibition (HI). Collected samples and associated laboratory results are summarized in Table S2 (Suppl. 2).

### GPS tracking

To study their movement ecology following the initial HPAIV outbreak, we deployed GPS-GSM tags on breeding adult gannets with chicks which showed no HPAIV symptoms on the Bass Rock (11^th^ to 13^th^ August 2022) and on Rouzic (23^rd^ to 26^th^ August 2022), about two months after the first signs of the disease were noticed in both colonies. GPS-GSM tags are pre-programmed to send data to a server using the 2G mobile-phone network so that data can be transmitted from anywhere within 2G coverage and accessed by the user in near-real-time. At Grassholm, birds were fitted with remotely-downloadable tags that send data via a UHF link to a base station deployed at the breeding colony in May 2022. At this location five tags were still operational 18^th^ August to 19^th^ September 2022.

For these GPS deployments, adults rearing chicks (2 to 8 weeks old) were caught at the nest and handled so as to minimize stress: the procedure lasted < 10 min and birds were kept in the shade. Tags were attached with Tesa® tape to the three central tail feathers. For Bass Rock gannets, we used nanofix tags (Pathtrack Ltd., solar powered, mass 18 g, 0.7% of bird body mass, dimensions 56×28×15mm plus ∼40 mm external whip antenna). These tags recorded one GPS position each 15 min or 30 min, depending on battery voltage. For Grassholm gannets, we used axy-trek remote tags (Technosmart solar powered, mass 15 g, < 0.5% of bird body mass, dimensions 52×25×15 mm). For Rouzic gannets, we used OrniTrack-9 tags (Ornitela, solar powered, mass 9 g, 0.3% of bird body mass, dimensions 37×19×12 mm, plus ∼10 cm external whip antenna). These tags were programmed to record a GPS position each 60 min at night, and each 15 min, 30 min or 60 min during the day, depending on battery voltage. We planned for tags to fall off once birds were starting to moult their tail feathers - previous deployments lasted up to 3 - 5 months (J. Jeglinski, D. Grémillet, unpublished data). At all three colonies we weighed gannets to the nearest 10 g. All birds were ringed with a metal ring and plastic ring with a unique alphanumeric combination. Once released, most birds returned immediately to their nests. We monitored chicks after capture until the captured parent had returned, generally immediately, at the latest within 1 h after capture.

### GPS Data processing

Data were analyzed using R version 4.1.2 (R Core Team 2021). Data from Rouzic, which are stored in the Movebank database, were imported with the *move* R package (Kranstauber et al., 2022) while data from Bass Rock and Grassholm were imported from csv files.

For the three datasets, at-sea trips were discriminated from periods in the colony when the birds were > 1 km away from the colony for > 1 h. Trips with time gaps longer than 10 h and incomplete trips were excluded, which represented 13.9% of trips for Bass Rock, 27.3% for Grassholm and 18.4% for Rouzic. At-sea locations were linearly interpolated with a 15min time resolution using the *pastecs* package (Grosjean et al., 2018). Two consecutive locations (= 30 min) in any other gannet colony were considered as prospecting.

We calculated trip characteristics (maximal distance to the colony in km, total trip duration in hrs and total distance travelled in km) as well as the time spent in the colony between at-sea trips in hrs, and present these averaged for each individual and then averaged by colony in Table 1. We fitted linear mixed models separately for each colony dataset using the respective trip characteristics for each bird as response variables to test whether log-transformed trip characteristics and time spent in the colony between at-sea trips changed significantly over time (converted as day since 10^th^ August 2022, first day of the study period). Bird ID was included as a random effect.

**Table 1.** Sample size, foraging trip characteristics, percentage of activities and colony attendance of northern gannets nesting on Rouzic, Grassholm and Bass Rock in 2022. Results are shown as mean ± SE

|  | Bass Rock | Grassholm | Rouzic |
| --- | --- | --- | --- |
| Total nb of equipped individuals | 10 | 12 | 15 |
| Nb of individuals retained | 10 | 5 | 13 |
| Total nb of foraging trips | 310 | 53 | 257 |
| Average nb of foraging trips per individual | 31 ± 4 | 10 ± 5 | 20 ± 3 |
| Total Nb of periods on the colony | 49 | 31 | 36 |
| Average nb of periods on the colony per individual | 30 ± 4 | 12 ± 5 | 18 ± 2 |
| Tracking period | 11/08/2022 -<br>12/10/2022 | 10/08/2022 –<br>18/09/2022 | 24/08/2022 –<br>28/09/2022 |
| Average tracking duration per individual (days) | 44.2 ± 5.1 | 13.8 ± 5.8 | 22.0 ± 2.4 |
| Maximal distance to the colony (km) | 98 ± 8 | 65 ± 8 | 62 ± 4 |
| Total distance travelled per trip (km) | 254 ± 21 | 191 ± 28 | 171 ± 11 |
| Trip duration (h) | 19 ± 2 | 16 ± 4 | 18 ± 1 |
| Time spent resting during a trip (%) | 21.7 ± 10 | 23.4 ± 3.2 | 42.4 ± 1.5 |
| Time spent foraging during a trip (%) | 54.4 ± 1.2 | 55.7 ± 2.9 | 36.9 ± 1.2 |
| Time spent flying during a trip (%) | 23.9 ± 0.8 | 20.9 ± 2.2 | 20.7 ± 0.9 |
| Time at the colony between at-sea trips (h) | 16 ± 1 | 18 ± 2 | 11 ± 1 |

### Inferring at-sea behavioural states

A 3-state hidden Markov model (HMM) was fitted to the at-sea interpolated location dataset from all three colonies combined, using the *moveHMM* package (Michelot et al., 2016). The states can be interpreted to reflect three different activities at sea: (1) resting, characterised by a small step length and low turning angle, (2) travelling, characterised by a long step length and low turning angle and (3) foraging, characterised by an intermediate step length and a large turning angle. A set of different initial parameter values was used to ensure that the global maximum log-likelihood had been reached. The Viterbi algorithm was used to classify the most likely behaviour at each location and proportions of each activity occurring during one at-sea trip were calculated, averaged by individual and then, by colony. We fitted logistic regressions to the proportions of each activity during a trip for each bird as response variables using the *glmmTMB* package (Brooks et al., 2017), accounting for zero-inflated data, to test whether activity proportions changed over time (converted as day since 10^th^ August 2022, first day of the study period). Bird ID was included as a random effect.

## Results

### Gannet colony dynamics

The impact of HPAIV on the Bass Rock was extremely high, with thousands of dead gannets in the colony and washing up on the surrounding beaches. A drone survey on 30^th^ June 2022 counted 21,277 gannets (Lane et al. unpubl. data), whereas a previous census counted 75,259 apparently occupied sites (AOS) in 2014 [38]. The two counts use slightly different metrics, but they suggest at least 71% decline in AOS during the 2022 HPAIV outbreak. Direct observations on Bass Rock also indicated that at least 75% of active nests disappeared during the 2022 breeding season (Lane et al. unpubl. data). On Grassholm, adult and chick mortality were extremely high with many thousands of birds dead on the colony and in the surrounding waters. On Rouzic, aerial pictures of the colony revealed that AOS numbers declined from 18,839 on 17/05/2022, to 8,625 on 28/08/2022 (54% decline during the breeding season). Observations via the live camera installed on the breeding colony estimated that >90% of chicks died during the same period (Suppl. 1). More detailed, weekly counts performed between 11/07/2022 and 30/09/2022 on five sections of the colony containing 40-102 AOS, suggested that between 58-87% of breeding adults died, and between 67-94% of the chicks (see Suppl. 3).

### Characterization of the HPAIV outbreak

#### Bass Rock

Analysis of samples from dead gannets by the Animal and Plant Health Agency (APHA), UK, confirmed an outbreak of HPAIV H5N1 (Scottish Seabird Centre, pers. comm.).

#### Grassholm

Following unusual gannet mortality in late July, four birds from 21^st^ July were confirmed as positive for HPAIV H5N1 by APHA.

#### Rouzic island

The first necropsy samples collected from a dead adult found on the colony on 11^th^ July 2022 tested positive for HPAIV. Laboratory analyses conducted on samples collected from a bird found dead on the colony on 30^th^ September 2022 confirmed the circulation of HPAIV H5N1 clade 2.3.4.4b in the colony. HPAIV H5 was also detected in live birds: 1/9 cloacal swabs from adults subsequently equipped with GPS, and 2/7 external wipes from juveniles (recently fledged and captured on land/at sea) tested positive for HPAIV H5. All the cloacal and tracheal swabs collected from these juveniles, as well as from 5 other juveniles were classified as negative or inconclusive, suggesting varying detection probabilities across sample types (see Suppl. 2). Antibodies against H5N1 were detected in 4/7 juveniles (including the 2 with RT-PCR-positive external swipes). These individuals also tested positive for antibodies against H5N5 and H5N8, and to a lesser degree, for antibodies against H5N3, although higher titers were systematically observed for H5N1 (Suppl. 2), suggesting cross-reactions with other serotypes. None of the samples tested positive for HPAIV H7.

### Movement ecology

All 10 GPS tags deployed on Bass Rock gannets sent GPS positions. On Grassholm, 5 out of 11 deployed tags provided data during the study period. For Rouzic, 13 out of 15 tagged individuals remained in the English Channel. One gannet, sampled at tag deployment, tested positive to HPAIV and never provided GPS positions, and another (tested negative at tag deployment) initiated migration after being equipped, moving toward West Africa without touching land until the tag stopped sending data on 16^th^ November 2022. (Table 1). Average gannet body mass was 3230±298g (Bass Rock), 3004±325g (Grassholm), and 2920±240g (Rouzic), which is in line with breeding gannet body mass in previous years (Le Bot et al., 2019).

While alive, none of the gannets came ashore anywhere outside of their respective breeding colonies during the GPS data transmission period. Overall, gannets from Bass Rock, Grassholm and Rouzic did not change their space use across the 4-8 weeks study period and foraged consistently in the same areas (Fig. 1). Birds from Rouzic did not change their at-sea trip characteristics over time (similar duration, maximal distance to the colony and total distance travelled; Table 2; Fig. 2a-c), but they slightly decreased the time spent at the colony between at-sea trips (Table 2; Fig. 2d). Among them, one individual was found dead on a beach on the island of Guernsey (82 km Northeast of Rouzic), 14 days after being equipped. This individual was not tested for HPAIV when found dead, and its behaviour for the time period during which we received GPS data did not differ from that of other Rouzic birds we tracked. Birds nesting on Bass Rock spent the same amount of time attending their colony but increased the duration of their at-sea trips marginally, travelling slightly further and longer in distance and time (Table 2; Fig. 2). Birds from Grassholm did not change any trip characteristics nor their nest attendance patterns. Considering their activity budgets during trips, birds maintained the same proportions of time spent foraging, flying and resting on the water, regardless of their colony (Table 2; Fig. 3). For the 3 colonies, compared to early chick-rearing in 2019, at-sea trips were 47-58% shorter in maximal distance and total distance travelled and 10-53% shorter in duration (Suppl. 4).

**Figure 1.**
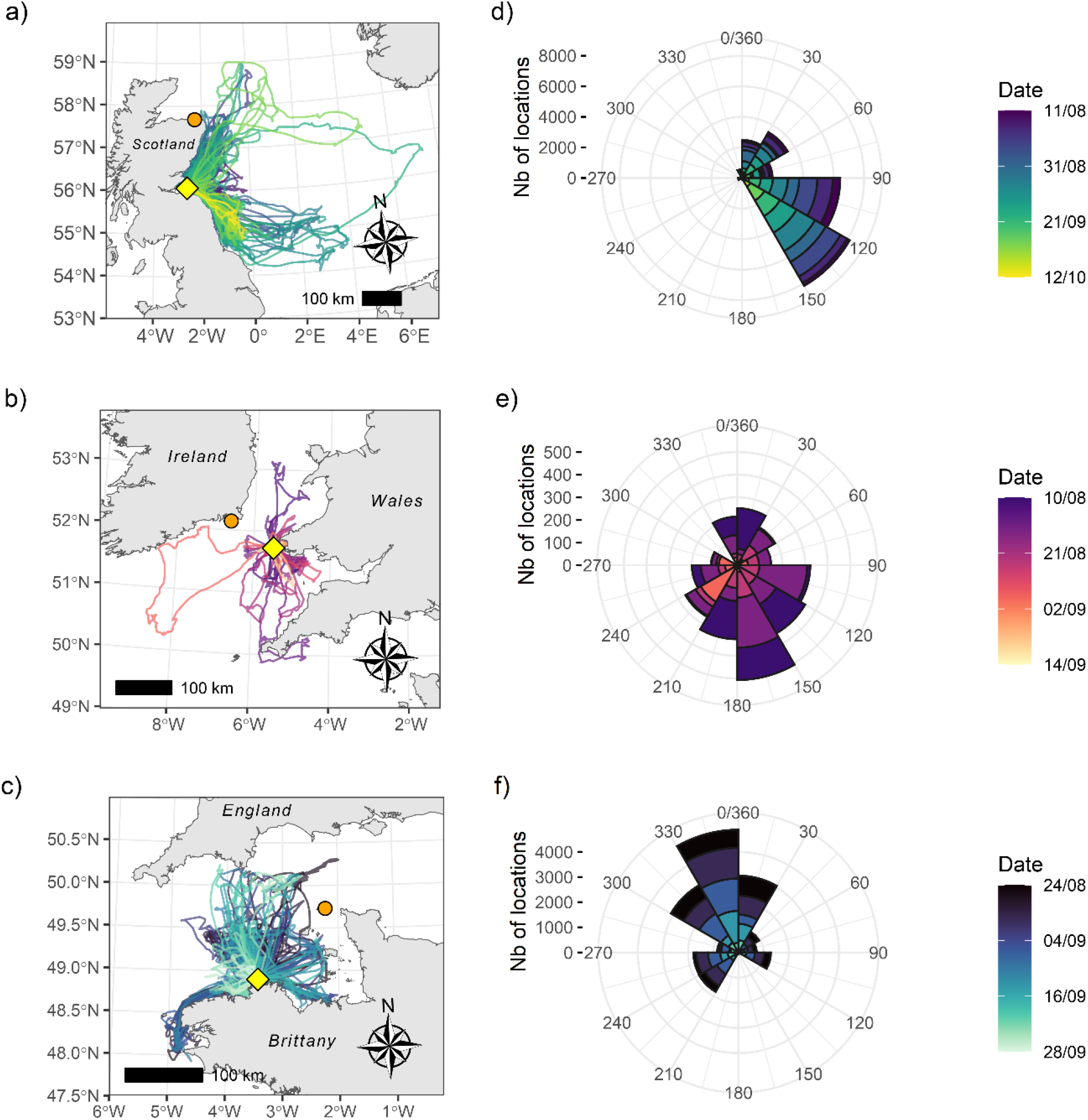
Tracks of northern gannets nesting in 2022 on (a) Bass Rock, Scotland, (b) Grassholm, Wales and (c) Rouzic, Brittany, (yellow diamonds). Orange circles represent the closest neighbouring gannet colony to the respective study colonies. Gradual changes in colour represent time in days since start of GPS tracking. (d-f) Rose diagrams illustrate the bearings of at-sea locations in 30 degree bins. The centre of each rose diagram represents the colony location, the length of each wedge represents the number of locations recorded in that direction over time, in coloured weekly bins.

**Table 2.** Results from the models testing the effects of time (dates between August and October 2022) on northern gannet trip characteristics and nest attendance (linear mixed models) and proportion of time spent in the 3 inferred behavioural states (logistic regressions). Individual identity is included as a random effect. Significant results (p < 0.05) are in bold.

|  | Bass Rock |  |  |  | Grassholm |  |  |  | Rouziec |  |  |  |
| --- | --- | --- | --- | --- | --- | --- | --- | --- | --- | --- | --- | --- |
|  | Estimate<br>± SE | df | t or z-<br>value | p-value | Estimate ±<br>SE | df | t or z-<br>value | p-value | Estimate ±<br>SE | df | t or z-<br>value | p-value |
| **Maximal distance** | **0.009 ±**<br>**0.005** | **299** | **2.03** | **0.043** | -0.012 ±<br>0.014 | 44 | -0.85 | 0.40 | -0.003 ±<br>0.006 | 243 | -0.53 | 0.60 |
| **Total distance**<br>**travelled** | **0.011 ±**<br>**0.005** | **299** | **2.34** | **0.02** | -0.017 ±<br>0.016 | 44 | -1.07 | 0.29 | $1.8e10^{-5} \pm$<br>$6.0e10^{-3}$ | 243 | 0.003 | 0.99 |
| **Trip duration** | **0.013 ±**<br>**0.004** | **299** | **3.29** | **0.001** | -0.014 ±<br>0.019 | 44 | -0.74 | 0.46 | 0.011 ±<br>0.006 | 243 | 1.73 | 0.09 |
| **Time at the colony**<br>**between at-sea trips** | -0.003 ±<br>0.003 | 288 | -0.77 | 0.45 | -0.011 ±<br>0.0010 | 56 | -1.13 | 0.26 | **-0.015 ±**<br>**0.007** | **220** | **-2.09** | **0.038** |
| **Proportion of a trip**<br>**spent resting** | 0.0053 ±<br>0.0086 | 307 | 0.63 | 0.53 | -0.05 ± 0.05 | 47 | -1.18 | 0.24 | 0.018 ±<br>0.014 | 253 | 1.24 | 0.21 |
| **Proportion of a trip**<br>**spent foraging** | -0.0017 ±<br>0.007 | 307 | -0.24 | 0.81 | 0.037 ±<br>0.034 | 47 | 1.05 | 0.29 | -0.014 ±<br>0.015 | 254 | -0.91 | 0.36 |
| **Proportion of a trip**<br>**spent flying** | -0.0028 ±<br>0.008 | 307 | -0.33 | 0.74 | -0.0021 ±<br>0.040 | 47 | -0.05 | 0.96 | -0.008 ±<br>0.018 | 254 | -0.43 | 0.67 |

**Figure 2.**
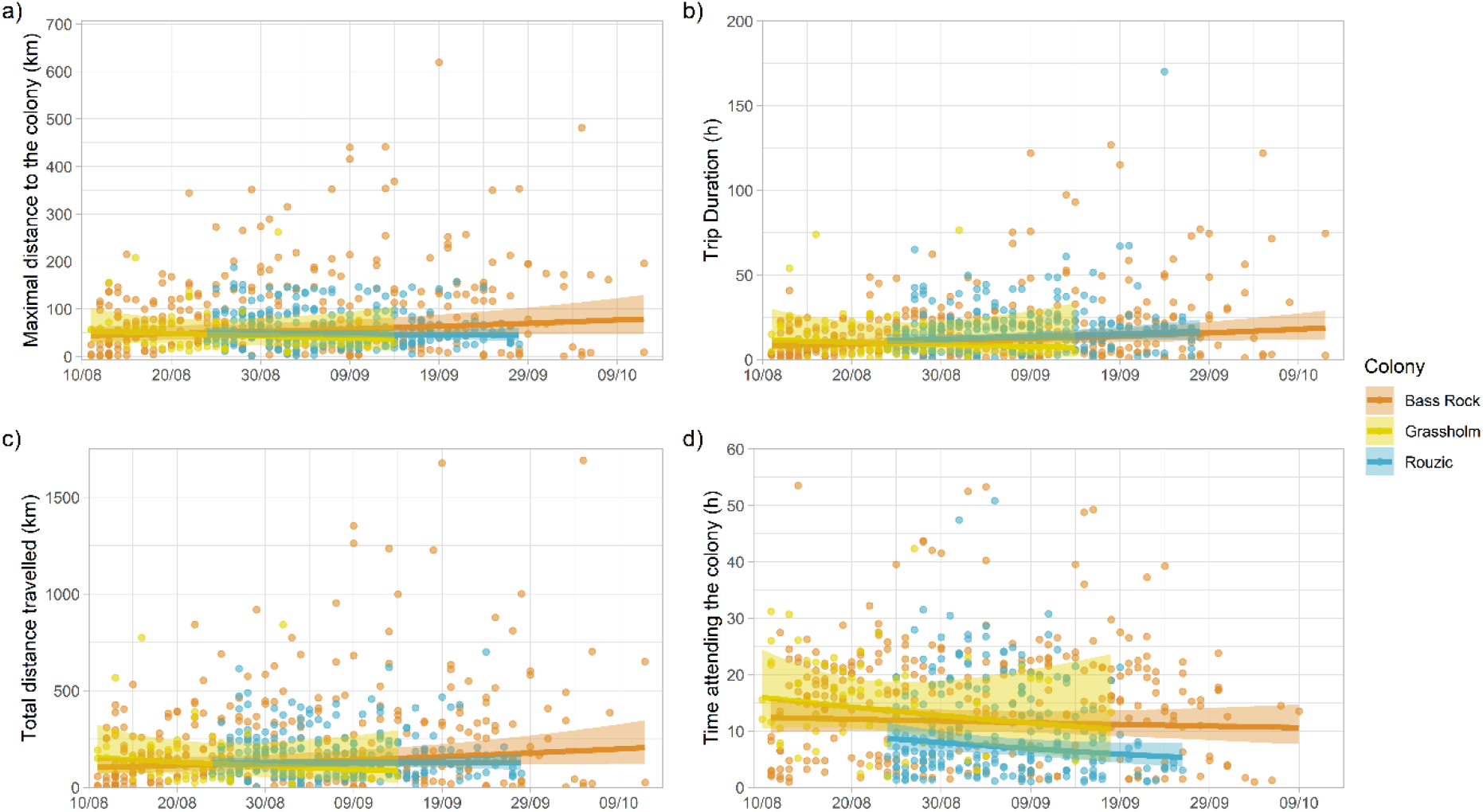
Trip characteristics and nest attendance of northern gannets nesting on Bass Rock (orange), Grassholm (yellow) and Rouzic (cyan), according to dates in August to October 2022. The dots represent raw data and the line and shadow the predicted average slope ± SE.

**Figure 3.**
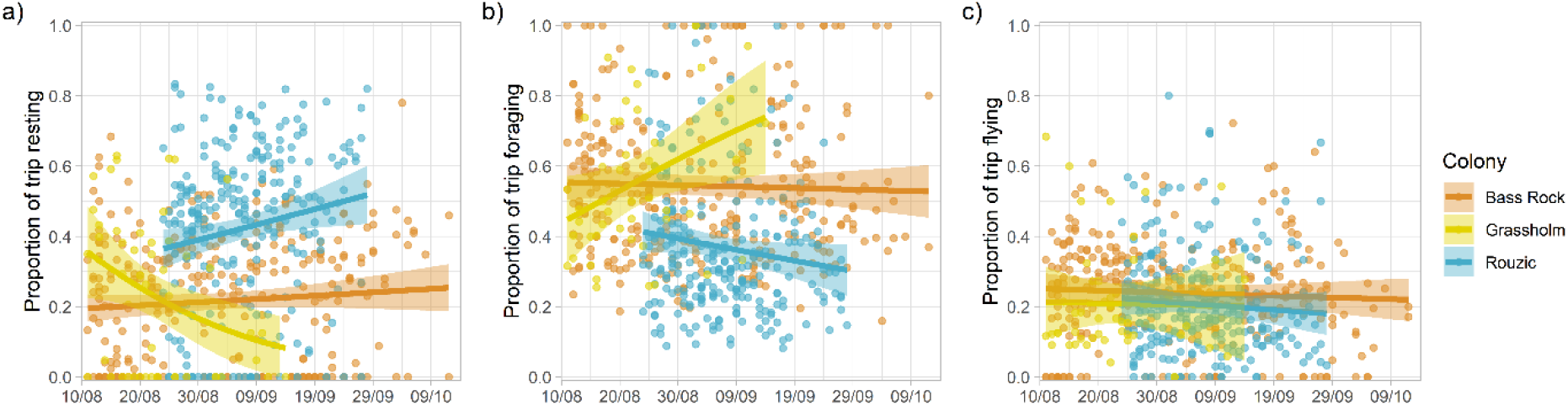
Proportion of inferred behavioural states [(a) resting, (b) foraging, (c) flying] during at-sea trips of northern gannets nesting on Bass Rock (orange), Grassholm (yellow) and Rouzic (cyan), according to dates in August to October 2022. The dots represent raw data and the line and shadow, predicted average slope ± SE.

## Discussion

### Epidemiology

Our analyses confirmed that northern gannets contracted HPAIV, and other investigations also pointed to HPAIV as the cause for 2022 seabird mass-mortalities in western Europe (Falchieri et al., 2022). More specifically, we identified H5N1 (clade 2.3.4.4b) in Rouzic birds. This contrasts with recent HPAIV outbreaks in southern African seabirds (Molini et al., 2020), which were caused by H5N8 (clade 2.3.4.4b). Our study adds to the evidence that HPAIV has become established in seabirds, and will probably recurrently re-emerge, as observed in southern Africa (Molini et al., 2020). We also found that some juvenile gannets had H5N1 antibodies, but no live virus (Suppl. 2). This is in line with recent investigations (Caliendo et al., 2022) suggesting that birds that survived HPAIV develop an immune response, which could protect against future infections. Further longitudinal monitoring will nonetheless be necessary to fully test this hypothesis. Those studies will also clarify the factors leading to intra and interspecific heterogeneity in avian responses to HPAIV (Keawcharoen et al., 2008), which are key to our understanding of the potential role of different species as HPAIV maintenance host or spreaders (Huang et al., 2019).

Only one of the 15 birds equipped with GPS on Rouzic tested HPAIV positive using swab samples (Suppl. 2), and gannets fitted with GPS on Bass Rock and Grassholm could not be tested. Thus, the epidemiological status of most tracked gannets was not known. Ideally, swab and plasma samples should have been collected in all birds. Yet, even in the presence of both sample types, some uncertainty would remain. Notably, a positive plasma sample could reflect a former, rather than an ongoing infection, and such samples are not adequate to infer whether a bird is a HPAIV carrier and potential spreader at the time of the blood sample. Indeed, experimentally infected birds stop excreting the virus after a week or less (van den Brand et al., 2018). Swab samples are therefore theoretically adequate during this brief excretion period, but half of those for gannets equipped with GPS on Rouzic were inconclusive. Overall, we may therefore conclude that HPAIV was extremely widespread and circulating at all three gannet colonies given observational data on the degree of mortality, and that an unknown proportion of GPS-tracked birds could have been symptomatic or asymptomatic carrier of the virus. This knowledge gap clearly calls for further epidemiological studies during future seabird breeding seasons.

### Movement ecology

We did not detect any change in breeding colony fidelity in the context of HPAIV outbreaks that started approximately two months previous to data collection. None of the tracked gannets came ashore outside of their respective breeding colonies while alive. We also found that GPS-tracked gannets did not significantly modify their foraging patterns across the study period (Table 2; Fig. 1-3). Since our visits to gannet breeding colonies were extremely limited due to the restrictions in place, we could not monitor the survival of the chicks of the GPS-tracked gannets. Some birds may have lost their chicks during the study period. Yet, all tracked birds continued to regularly commute between at-sea feeding sites and their respective colonies, and none of them rested on the mainland or on other islands in-between trips.

Further, gannet foraging effort seemed lower than during the 2019 breeding season (hence before HPAIV) at our three study sites (Grémillet et al., 2006; Hamer et al., 2007; Patrick et al., 2014) (Suppl. 4). Therefore, gannets surviving HPAIV may have benefited from relaxed density-dependence. Our data are not directly comparable with those collected in 2019, because we studied the late chick-rearing period of the birds, whereas most previous studies focused on early chick-rearing. Yet, late chick-rearing requires twice more energy than early chick-rearing (Montevecchi et al., 1984), and lower foraging effort during late chick-rearing in 2022 does point to reduced intra- and interspecific competition for food.

Our results are coherent with high breeding and foraging site fidelity in colonial seabirds (Danckwerts et al., 2021), and contrary to the idea that gannets that have witnessed and survived mass mortality at their breeding colony modify their movement patterns as a consequence of the disease outbreak and maintain these changes several months after the event. Thus, it appears that the capacity for long-term enhanced HPAIV transmission throughout the metapopulation and potentially throughout the co-existing seabird populations is low for adult breeding gannets. This is an essential piece of information that contributes to a better understanding of HPAIV transmission dynamics, since wild birds carrying HPAIV are recurrently presented as virus spreaders to farm poultry (H5N8 and Viruses, 2016, p. 8).

However, the data we presented here are collected following the initial disease outbreak to study long-term behavioural responses, and they are only representative of the adult breeding age class. Short-term responses of breeding birds, at the start of a disease outbreak, might differ from the behavioural and spatial consistency we demonstrated here. It might also be that age classes differ in their propensity to act as a vector for HPAIV. Immature gannets have been shown to perform visits to other breeding colonies throughout the breeding season (Votier et al., 2011) and thus likely regularly move between colonies. These prospecting movements would appear to require limited time/energy investment: For gannets from Bass Rock, the closest alternative colony is Troup Head on the Scottish mainland (ca. 210 km), for Grassholm it is Great Saltee in Ireland (ca. 90 km), and for Rouzic it is Ortac in the British Channel Islands (ca. 130 km) (Fig.1). Reaching these neighbouring colonies would take 3.2 hrs (Bass Rock), 1.4 hrs (Grassholm), or 2 hrs (Rouzic), assuming a 65 km.h^-1^ travelling speed (Grémillet et al., 2004). Forthcoming studies will clarify the respective potential of juvenile and immature gannets to act as vectors in HPAIV transmission. Also, even if rated as rare (Boulinier et al., 2016), at-sea HPAIV transmission may occur within, and between seabird species. This might be facilitated by aggregations at natural foraging sites or whilst feeding on fishery wastes behind trawlers, or through kleptoparasitism and scavenging (Wille et al., 2016). Studying interspecific fine scale space-use overlap at sea is therefore a worthwhile avenue for future monitoring and evaluation of disease transmission scenarios (Boulinier, 2023).

### Outlook

It remains unclear how gannets contracted HPAIV, since they were either at sea, or at their breeding colony during the outbreak. Larids (*Larus sp*.) or great skuas that klepto-parasitise gannets might be potential vectors since they use similar terrestrial and marine habitats (Falchieri et al., 2022), but further analyses will be necessary to test these hypotheses. HPAIV first affected gannets in Iceland, Québec and Scotland (from early April 2022), and subsequently arrived in Brittany (from early July 2022). Hence, it is possible that gannets moving between these colonies may have spread the virus, but this pathway is still hypothetical. Equally unknown are HPAIV dynamics during gannet migration, which started in September-October 2022 and lasted until February 2023. These inter-breeding movements take gannets from the British Isles and Brittany to the Mediterranean or West Africa (Fort et al., 2012). Off Africa, gannets meet with some of their North American conspecifics, as well as with a vast community of North Atlantic seabirds spending the winter in tropical waters (Grecian et al., 2016).

Our study is a key example of the common fate of wildlife and humanity facing global changes (Díaz et al., 2019). Gannets are affected by the direct and indirect consequences of climate change (heat waves and storms, changes in the distribution and abundance of marine resources), by pollutions (e.g. mercury in fish affecting seabird and human health), and by uncontrolled fishing operations which threaten food security for seabird and human populations alike (Grémillet et al., 2020). In addition, the current HPAIV outbreak demonstrates that the anthropogenic environmental crisis enhances the potential for emerging infectious disease, contaminating wild animals and humans (Carlson et al., 2022; Kuiken and Cromie, 2022). Indeed, current industrial farming practices are responsible for the emergence and the spread of HPAIV in poultry and subsequently in wild birds, potentially also in humans (Yamaji et al., 2020).

## Declarations

## Acknowledgements

For the Bass Rock fieldwork, we thank Sir Hew Dalrymple for the permission to conduct our research project. We also thank Maggie Sheddan, Jack Dale, John McCarter and the Scottish Seabird Centre for logistical support. For the Grassholm fieldwork we thank Dale Sailing. Field studies on Rouzic greatly benefited from the help of all staff at the Station Ornithologique de l’Ile Grande. We are also grateful for the help of Manon Amiguet, Yann Villaggi (DDPP22), Gauthier Poiriez & Paco Bustamante (LIENSs La Rochelle), and for advice provided by Grégoire Kuntz.

## Author contributions

DG, PP and SCV designed the study, with advice from SW and JJ. DG, PP, APr, TP, JJ, SCV, JM, AB, JF, NC, RJ, LL, AT, NS and JL were leading the field studies including funding acquisition, permit acquisition and field data collection. APo, PP, MA-Z, AG, NC and DG analysed the data. DG, APo, AG and JJ wrote the paper, with inputs from all other authors.

## Ethics

All procedures followed the *guidelines for the treatment of animals in behavioural research and teaching* of the *Association for the Study of Animal Behaviour*. Bass Rock: fieldwork was performed in partnership with the Scottish Seabird Centre with permission of the landowner of the Bass Rock, Sir Hew Dalrymple, and under a special method endorsement and ringing permit of the British Trust for Ornithology BTO to Jana Jeglinski. Colour ringing was performed under a permit to Jude Lane from the BTO. We conducted our fieldwork under an exemption to the general ringing ban in seabird colonies granted by NatureScot which permitted us to capture and tag gannets during the HPAIV outbreak. All fieldwork was performed with the required risks assessments and ethical approval in place. Grassholm: All research was carried out under licence (RSPB, Natural Resources Wales (#S091127-1), British Trust for Ornithology (BTO: A4257), the BTO Special Methods Panel and the UK Home Office (30/3065). Rouzic: fieldwork operated under the Programme Personnel de baguage n°536 CRBPO/MNHN validation 2020-2024.

## Data availability

Data are available on the following archives: https://www.movebank.org/cms/webapp?gwt_fragment=page=studies,path=study2296851236 ; https://datadryad.org/stash/share/MmNkFWcQV79wnYDMQ8VTav-jRmHBUomyRohKVoAsNvE and https://www.movebank.org/cms/webapp?gwt_fragment=page=studies,path=study2658117564. All statistical analyses can be found on the GitHub repository (https://github.com/auponchon/NorthernGannets2022).

