## SupplementaryMaterial1 for "Strong breeding colony fidelity in northern gannets following High Pathogenicity Avian Influenza Virus (HPAIV) outbreak"

**Supplementary material 1:**

Examples of how the 2022 HPAIV outbreak impacted the Bass Rock (a,b & c) and Rouzic (d,e &f) northern gannet colonies. Timing of the outbreak differed between colonies and between colony sections (see results in the main document, and Suppl. Mat. 3).

Note that the top picture of Bass Rock is from July 2020 for reference.

**a) Bass Rock - 22$^{nd}$ July 2020, ~906 adult gannets**

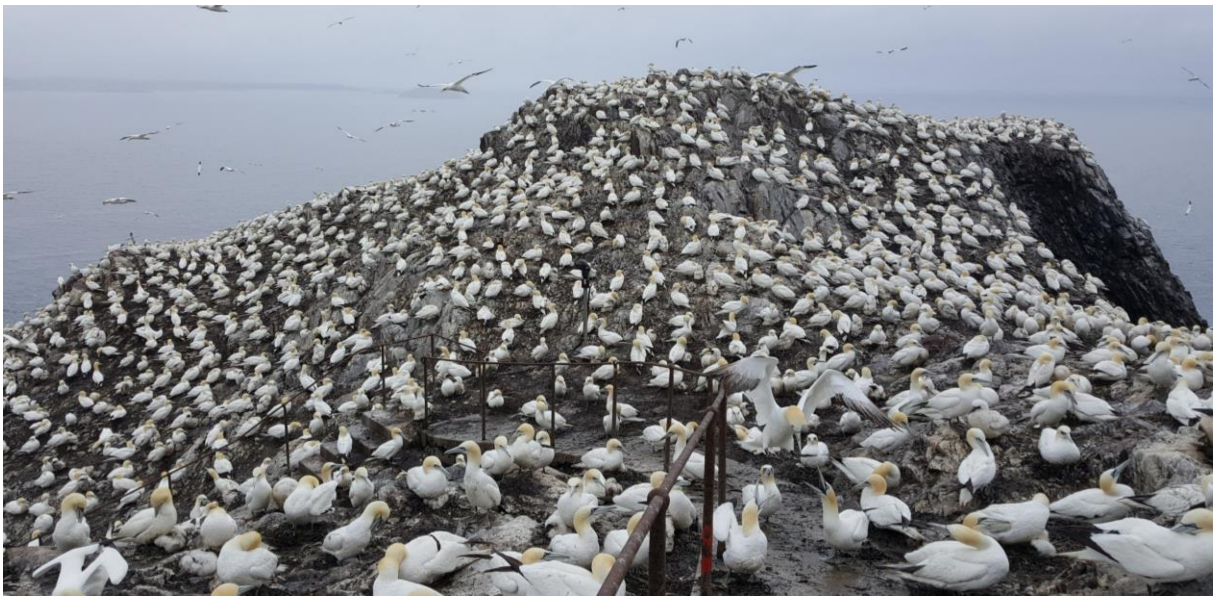

**b) Bass Rock - 21$^{st}$ June 2022, 326 adult gannets (alive)**

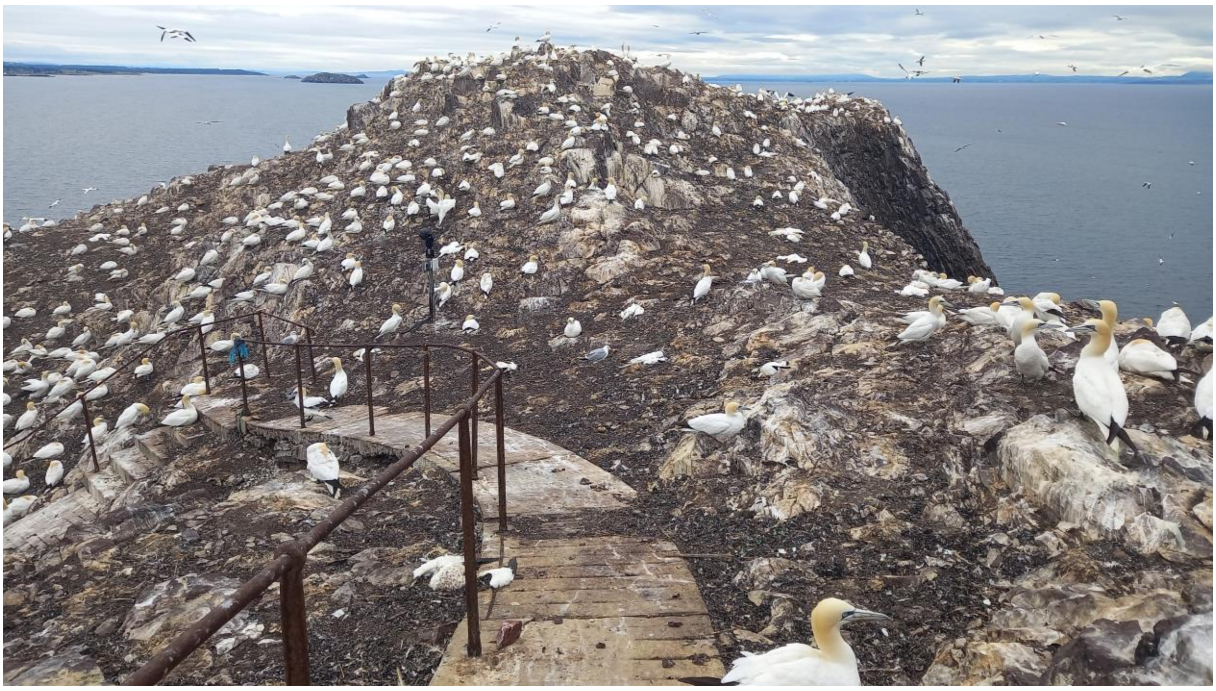

**c) Bass Rock - 30$^{th}$ June 2022, 87 adult gannets (alive)**

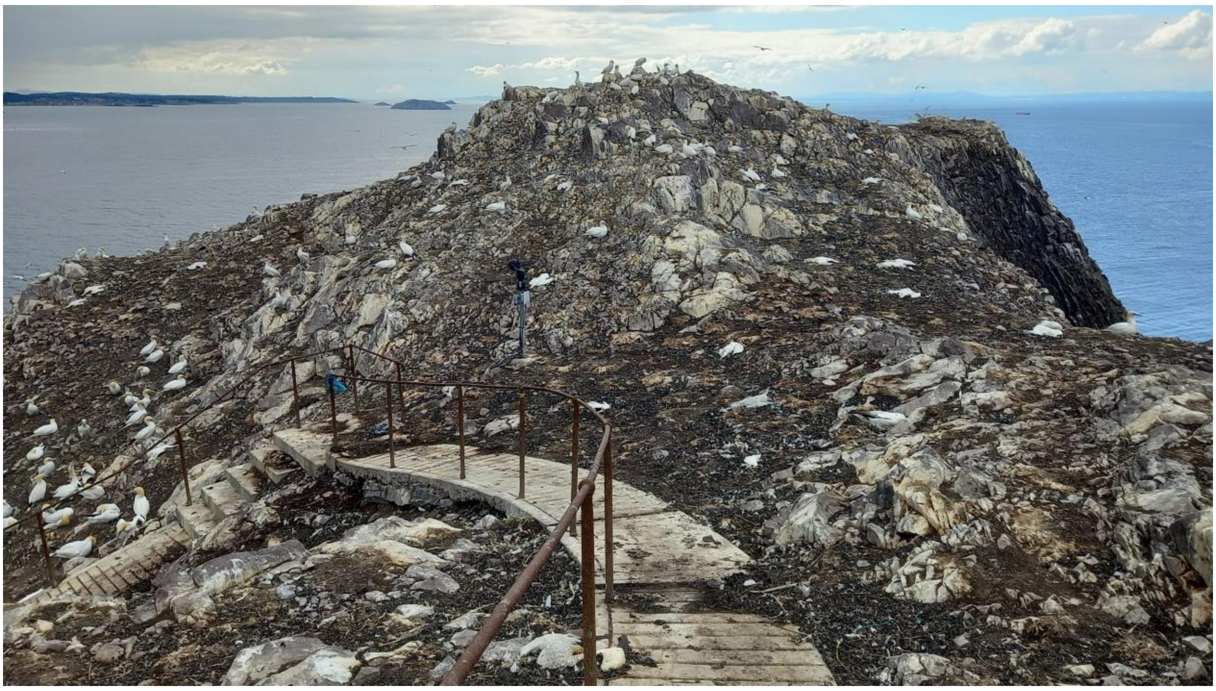

**Picture credits: Jude Lane, RSPB**

**d) Rouzic - 2nd August 2022, 718 gannets (alive)**

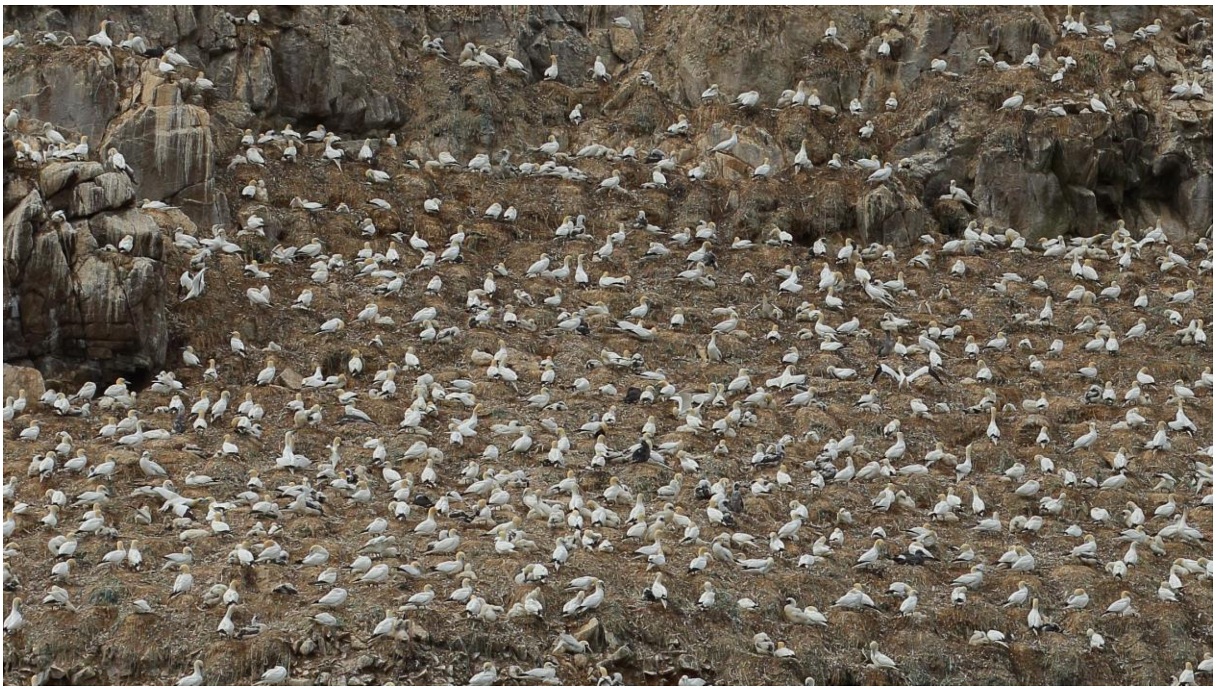

**e) Rouzic - 23rd August 2022, 249 gannets (alive)**

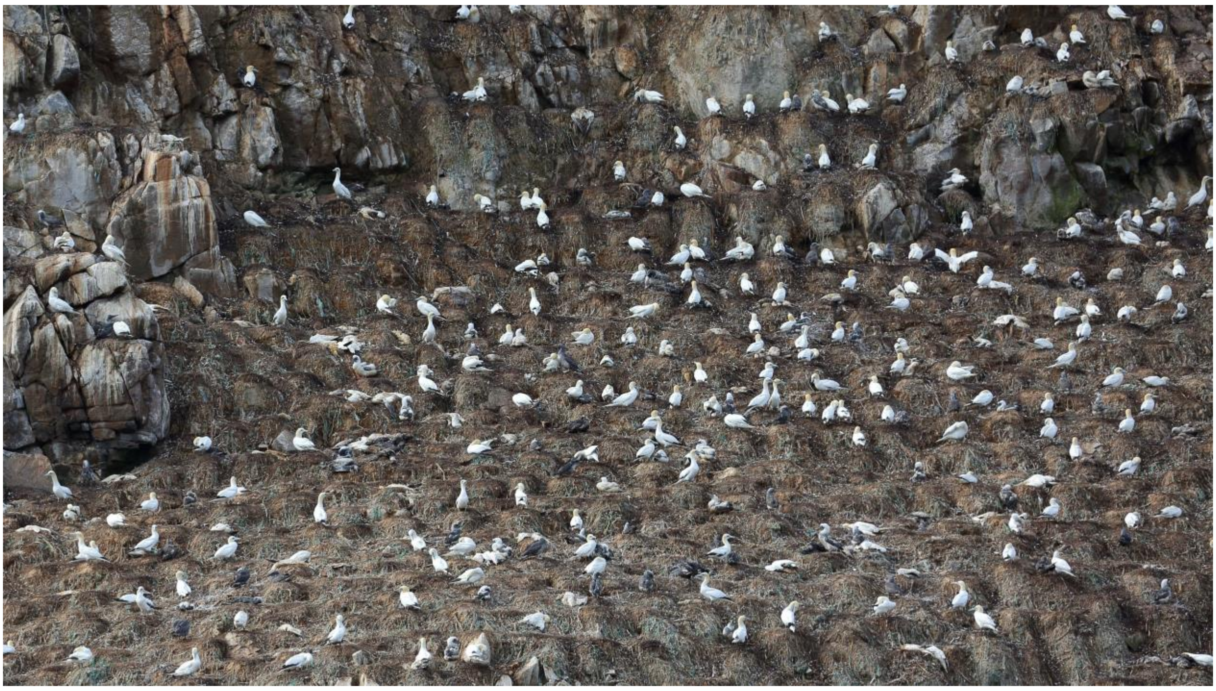

**f) Rouzic - 15th September 2022, 49 gannets (alive)**

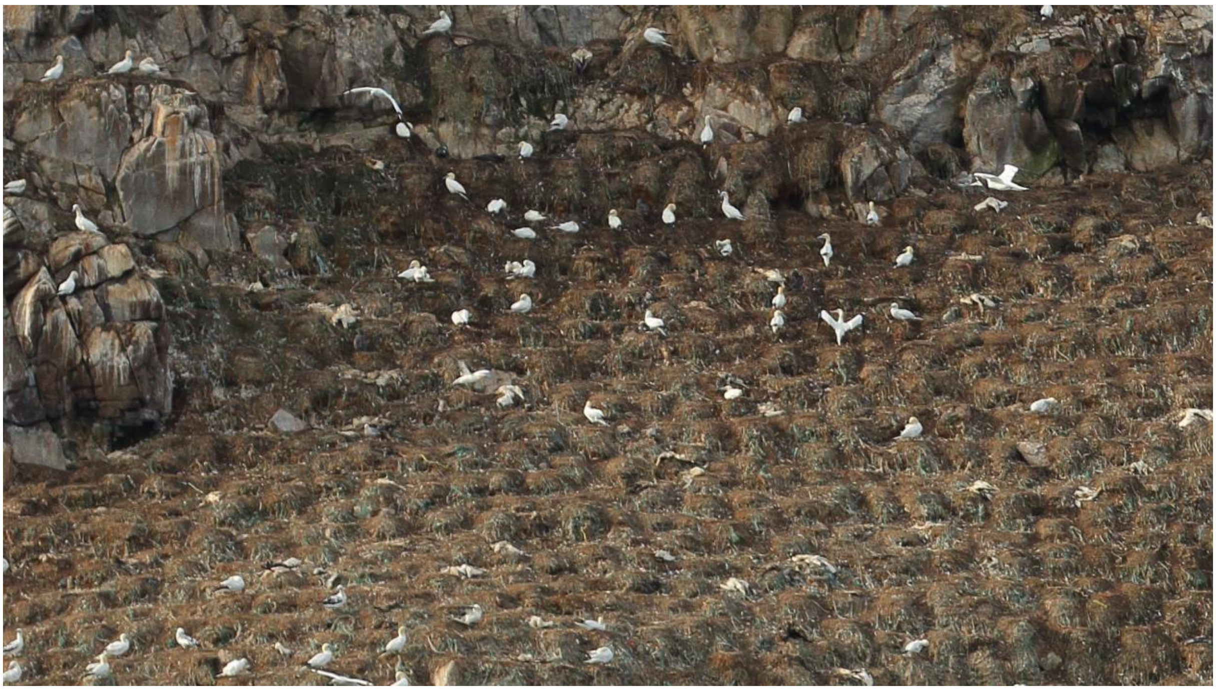

**Picture credits: Armel Deniau**
