## SupplementaryMaterial2 for "Strong breeding colony fidelity in northern gannets following High Pathogenicity Avian Influenza Virus (HPAIV) outbreak"

***Table S2.*** *Summary of the AIV diagnostic results for Northern gannets from Rouzic and surroundings. Tissue samples, swabs and wipes were analyzed by RT-PCR and classified as inconclusive (Inc.; host material genetic not detected), negative (Neg.; AIV M not detected) or positive (Pos.; AIV M, H5 and/or N1 detected). Plasma samples were analyzed by HI; results are reported as antibody titers (corresponding to the last positive dilution). Evidence of past or present infection with AIV are indicated in bold. The asterisk (*) indicates detection of H5 clade 2.3.4.4b by RT-PCR. Hashtags (#) indicate samples pooled across 2 individuals.*

| Sampling date | Sampling location | Bird age | Bird status | Comment | Lungs and/or trachea  RT-PCR | Cloacal swab  RT-PCR | Tracheal swab  RT-PCR | External wipe  RT-PCR | Plasma  HI | | | |
| --- | --- | --- | --- | --- | --- | --- | --- | --- | --- | --- | --- | --- |
|  |  |  |  |  |  |  |  |  | H5N3 | H5N1 | H5N5 | H5N8 |
| 23-25/08/2022 | Colony | Adult | Live | GPS-GSM |  | Neg. (M) |  |  |  |  |  |  |
| 23-25/08/2022 | Colony | Adult | Live | GPS-GSM |  | Neg. (M) |  |  |  |  |  |  |
| 23-25/08/2022 | Colony | Adult | Live | GPS-GSM |  | Inc. |  |  |  |  |  |  |
| 23-25/08/2022 | Colony | Adult | Live | GPS-GSM |  | Neg. (M) |  |  |  |  |  |  |
| 23-25/08/2022 | Colony | Adult | Live | GPS-GSM |  | Neg. (M) |  |  |  |  |  |  |
| 23-25/08/2022 | Colony | Adult | Live | GPS-GSM |  | Neg. (M) |  |  |  |  |  |  |
| 23-25/08/2022 | Colony | Adult | Live | GPS-GSM |  | Neg. (M) |  |  |  |  |  |  |
| 23-25/08/2022 | Colony | Adult | Live | GPS-GSM |  | Inc. |  |  |  |  |  |  |
| 23-25/08/2022 | Colony | Adult | Live | GPS-GSM |  | Inc. |  |  |  |  |  |  |
| 23-25/08/2022 | Colony | Adult | Live | GPS-GSM |  | Inc. |  |  |  |  |  |  |
| 23-25/08/2022 | Colony | Adult | Live | GPS-GSM |  | Inc. |  |  |  |  |  |  |
| 23-25/08/2022 | Colony | Adult | Live | GPS-GSM |  | Inc. |  |  |  |  |  |  |
| 23-25/08/2022 | Colony | Adult | Live | GPS-GSM |  | Neg. (M) |  |  |  |  |  |  |
| 23-25/08/2022 | Colony | Adult | Live | GPS-GSM |  | Neg. (M) |  |  |  |  |  |  |
| 23-25/08/2022 | Colony | Adult | Live | GPS-GSM |  | **Pos. (M, H5)** |  |  |  |  |  |  |
| 20/09/2022 | Sea | Juvenile | Live |  |  | Inc. or Neg.^#^ | Inc.^#^ |  |  |  |  |  |
| 20/09/2022 | Sea | Juvenile | Live |  |  | Inc. or Neg.^#^ | Inc.^#^ |  |  |  |  |  |
| 20/09/2022 | Sea | Juvenile | Live |  |  | Inc. or Neg.^#^ | Inc. or Neg.^#^ |  |  |  |  |  |
| 20/09/2022 | Sea | Juvenile | Live |  |  | Inc. or Neg.^#^ | Inc. or Neg.^#^ |  |  |  |  |  |
| 20/09/2022 | Sea | Juvenile | Live |  |  | Neg. | Inc. |  |  |  |  |  |
| 30/09/2022 | Colony | Adult | Dead |  | **Pos. (M, H5*, N1)** |  |  |  |  |  |  |  |
| 30/09/2022 | Colony | Adult | Dead |  | Inc. |  |  |  |  |  |  |  |
| 30/09/2022 | Colony | Juvenile | Dead |  | Inc. |  |  |  |  |  |  |  |
| 30/09/2022 | Colony | Juvenile | Dead |  | Inc. |  |  |  |  |  |  |  |
| 30/09/2022 | Colony | Juvenile | Live |  |  | Inc. or Neg.^#^ | Inc.^#^ | Neg. | <4 | <2 | <2 | <2 |
| 30/09/2022 | Colony | Juvenile | Live |  |  | Inc. or Neg.^#^ | Inc.^#^ | Neg. | <4 | <2 | <2 | <2 |
| 30/09/2022 | Colony | Juvenile | Live |  |  | Inc. or Neg.^#^ | Inc.^#^ | **Pos. (M, H5)** | <4 | **128** | **64** | **64** |
| 30/09/2022 | Colony | Juvenile | Live |  |  | Inc. or Neg.^#^ | Inc.^#^ | **Pos. (M, H5)** | 4 | **64** | **32** | **16** |
| 30/09/2022 | Colony | Juvenile | Live |  |  | Inc. or Neg.^#^ | Inc.^#^ | Neg. | <4 | <2 | <2 | <2 |
| 30/09/2022 | Colony | Juvenile | Live |  |  | Inc. or Neg.^#^ | Inc.^#^ | Neg. | <4 | **128** | **128** | **128** |
| 30/09/2022 | Colony | Juvenile | Live |  |  | Neg. | Inc. | Neg. | 8 | **128** | **32** | **32** |
