## SupplementaryMaterial3 for "Strong breeding colony fidelity in northern gannets following High Pathogenicity Avian Influenza Virus (HPAIV) outbreak"

Supplementary material 3: **Detailed analysis of nest failures at the Rouzic Northern gannet breeding colony, Sept-Iles Archipelago, Brittany, France.**

Methods: 40 to 102 apparently occupied sites (AOS) were selected at random within five different areas of the Rouzic colony (see pictures S3.1-S3.5 below). Between July-September 2022, pictures of those areas/nests were taken from the sea on nine dates for four areas (one per week), and on 14 dates with a remote CCTV installed on the colony, for the fifth area. Using these pictures, we determined the contents of each AOS, using the following categories: (1) Adult; (2) Egg; (3) small chick (< 2 weeks of age); (4) chick between 2 and 12 weeks of age; (5) juvenile about to fledge (chick > 12 weeks of age).

Using AOS contents observed during the first round of observations in July, before HPAIV significantly affected northern gannets and their breeding performance, we calculated potential maximum chick production per area, P_Pmax_, as the total number of observed chicks and eggs, divided by the total number of AOS for this area. For this initial period, we calculated potential minimum chick production per area, P_Pmin_, as the total number of observed chicks older than two weeks (and hence likely to survive), divided by the total number of AOS for this area.

During later observations, when HPAIV was affecting the colony at a time when chicks should be larger and about to fledge in August/September, we calculated actual maximum chick production per area, P_Amax_, as the total number of observed chicks older than 12 weeks, divided by the total number of AOS for this area. We had doubt on the fate of some chicks and then calculated actual minimum chick production per area, P_Amin_, as the total number of observed chicks older than 12 weeks for which there was no doubt about their survival, divided by the total number of AOS for this area.

To assess the impact of HPAIV on chick production, we calculated the range of decrease in chick production for each area as following:

$$Minimum chick loss=\frac{P_{Amax}-P_{Pmin}}{P_{Pmin}}\times100$$

$$Maximum chick loss=\frac{P_{Amin}-P_{Pmax}}{P_{Pmax}}\times100$$

In addition, we assessed the impact of HPAIV on adult survival. We evaluated the range of living adults in each AOS at the end of September based on the following criteria: (0) No adult was observed on AOS after the first cases of HPAIV, (0-1) One adult observed on AOS after the first cases of HPAIV at one date, (1) One adult observed on AOS after the first cases of HPAIV at several dates, (0-2) the pair was observed on AOS after the first cases of HPAIV at one date or a fledging chick was observed at the beginning of fledging period plus no adult was observed on AOS after the first cases of HPAIV, (1-2) a fledging chick was observed at the beginning of fledging period plus one adult was observed on AOS after the first cases of HPAIV at several dates, (2) one to two adults were observed on AOS after the first cases of HPAIV at several dates or a fledging chick was observed at the end of fledging period. We calculated the maximum number of living adults, S_max_, and the minimum number of living adults, S_min_, for each area. Considering that the expected number of living adults without the HPAIV event, N, is the total number of AOS multiplied by 2, we deduced the decrease in adult survival as follows:

$$Minimum adult loss=\frac{S_{max}-N}{N}\times100$$

$$Maximum adult loss=\frac{S_{min}-N}{N}\times100$$

Results: HPAIV greatly impacted the Northern gannet breeding site of Rouzic, with potentially 94.5% of all AOSs experiencing demographic consequences (Table S3.1). The potential chick production estimated at 42.2% at the beginning of breeding period decreased to 10.5% with the HPAIV outbreak (Table S3.1). The loss of chick production ranged between -67.3% (± SD: 24.2%) and -71.8% (± 26.4%), with a maximum local decrease of -94.3% in the West area (Table S3.1). We also found that HPAIV strongly affected adult survival, with an estimated adult loss ranging between -58.3% (± 13.9%) and -86.9% (± 4.7%), potentially representing the death of 19,514 to 27,187 adult gannets on Rouzic (Table S3.1). Note that chick survival varied more than adult survival between the five areas.

Table S3.1. Summary for the apparently occupied nest (AOS) monitoring at five areas on Rouzic Island and for the loss of chicks and adults. East, Northeast, Northwest and West areas were monitored by sea on nine occasions: 11/07, 27/07, 02/08, 12/08, 23/08, 07/09, 15/09, 20/09, 30/09. Plateau area was monitored by remote CCTV on 14 occasions (05/07, 11/07, 18/07, 25/07, 28/07, 01/08, 08/08, 16/08, 23/08, 30/08, 05/09, 12/09, 20/09).

|  | East | Northeast | Northwest | West | Plateau |
| --- | --- | --- | --- | --- | --- |
| Number of AOS monitored | 40 | 40 | 40 | 60 | 102 |
| Maximum number of AOS in May | 1567 | 2606 | 2268 | 5573 | 3549 |
| First confirmed case of HPAIV | Start September | Mid-August | Start September | Start August | Start July |
| Number of AOS affected by HPAIV | 57.5% to 90.0% | 87.5% to 97.5% | 42.5% to 95.0% | 42.5% to 95.0% | 74.5% to 95.1% |
| Potential chick production, P_Pmin_ to P_Pmax_ | 11 ; 11 | 23 ; 24 | 10 ;10 | 33 ; 35 | 38 ; 50 |
| Actual chick production, P_Amin_ to P_Amax_ | 4 ; 4 | 5 ; 8 | 7 ; 7 | 2 ; 3 | 4 ; 5 |
| Chick production before HPAIV | 27.5% | 57.5% to 60% | 25.0% | 55.0% to 58.3% | 37.3% to 49.0% |
| Chick production during HPAIV | 10.0% | 12.5% to 20% | 17.5% | 3.3% to 5.0% | 3.9% to 4.9% |
| Minimum chick production loss | -63.6% | -65.2% | -30.0% | -90.9% | -86.8% |
| Maximum chick production loss | -63.6% | -79.2% | -30.0% | -94.3% | -92.0% |
| Adult survival, S_min_ to S_max_ | 13 ; 46 | 6 ; 25 | 10 ; 44 | 12 ; 32 | 39 ; 78 |
| Minimum adult loss | -42.5% | -68.8% | -45.0% | -73.3% | -61.2% |
| Maximum adult loss | -83.8% | -92.5% | -87.5% | -90.0% | -80.9% |

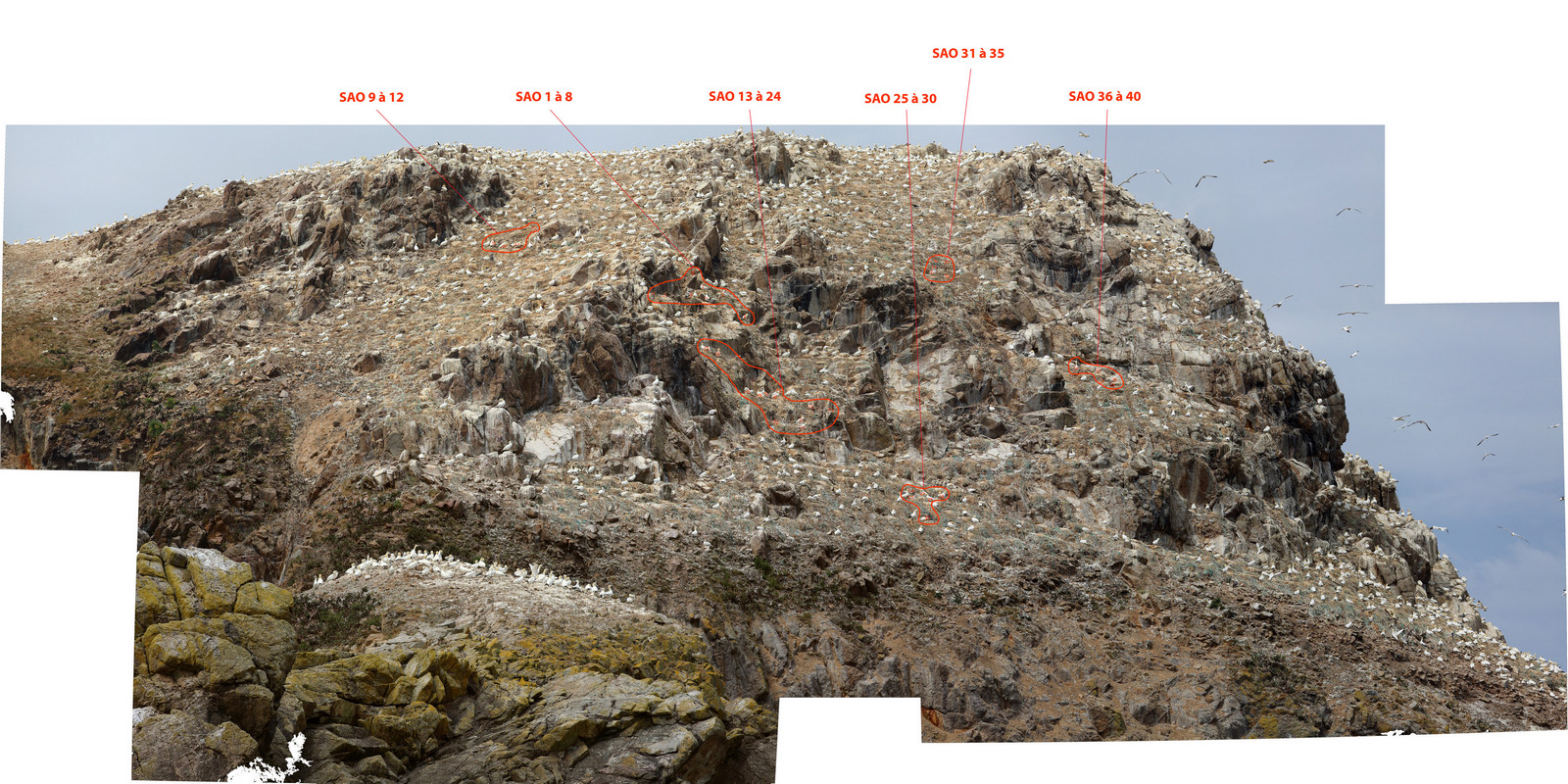

Picture S3.1. East area with the 40 AOS followed indicated in red.

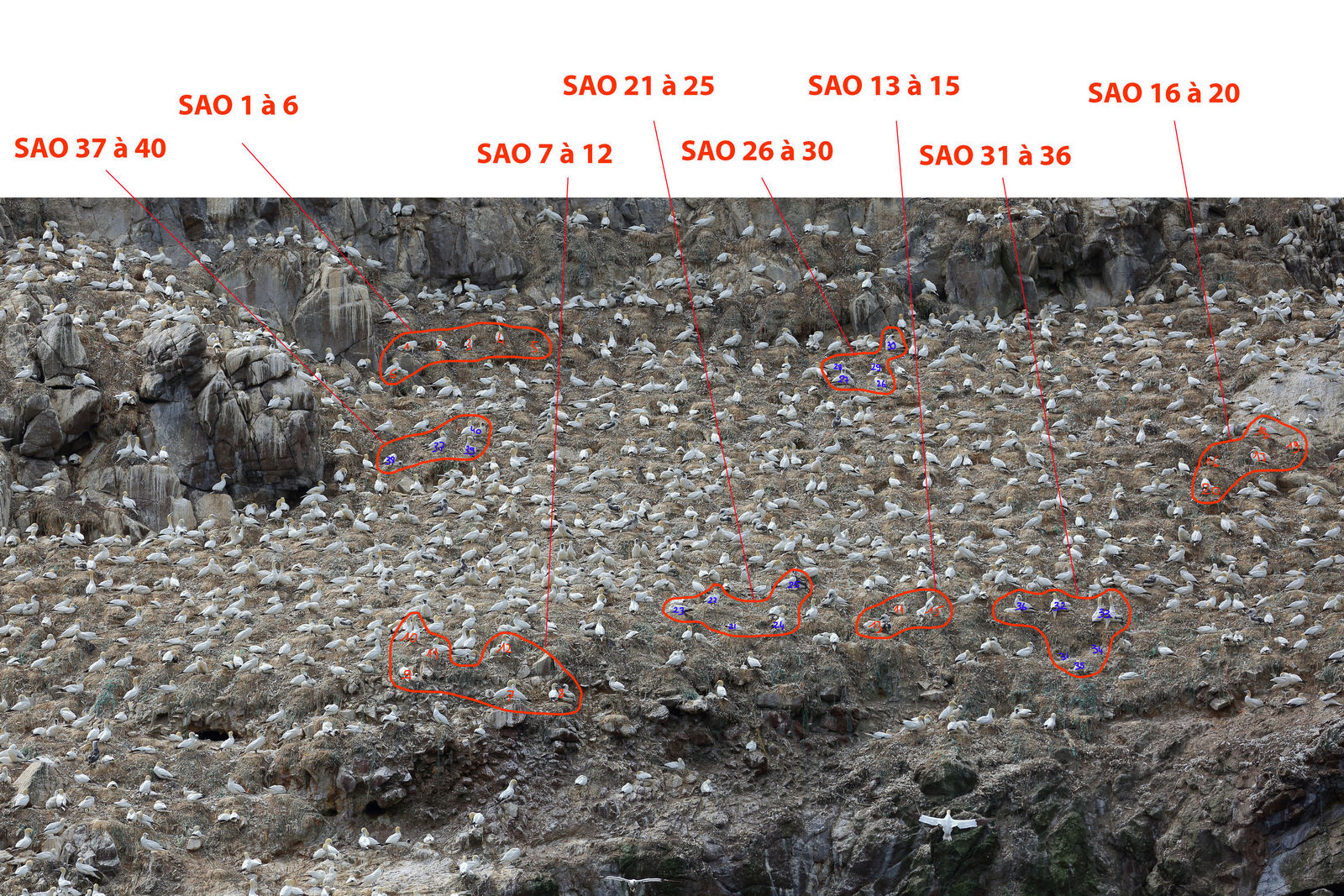

Picture S3.2. Northeast area with the 40 AOS followed indicated in red.

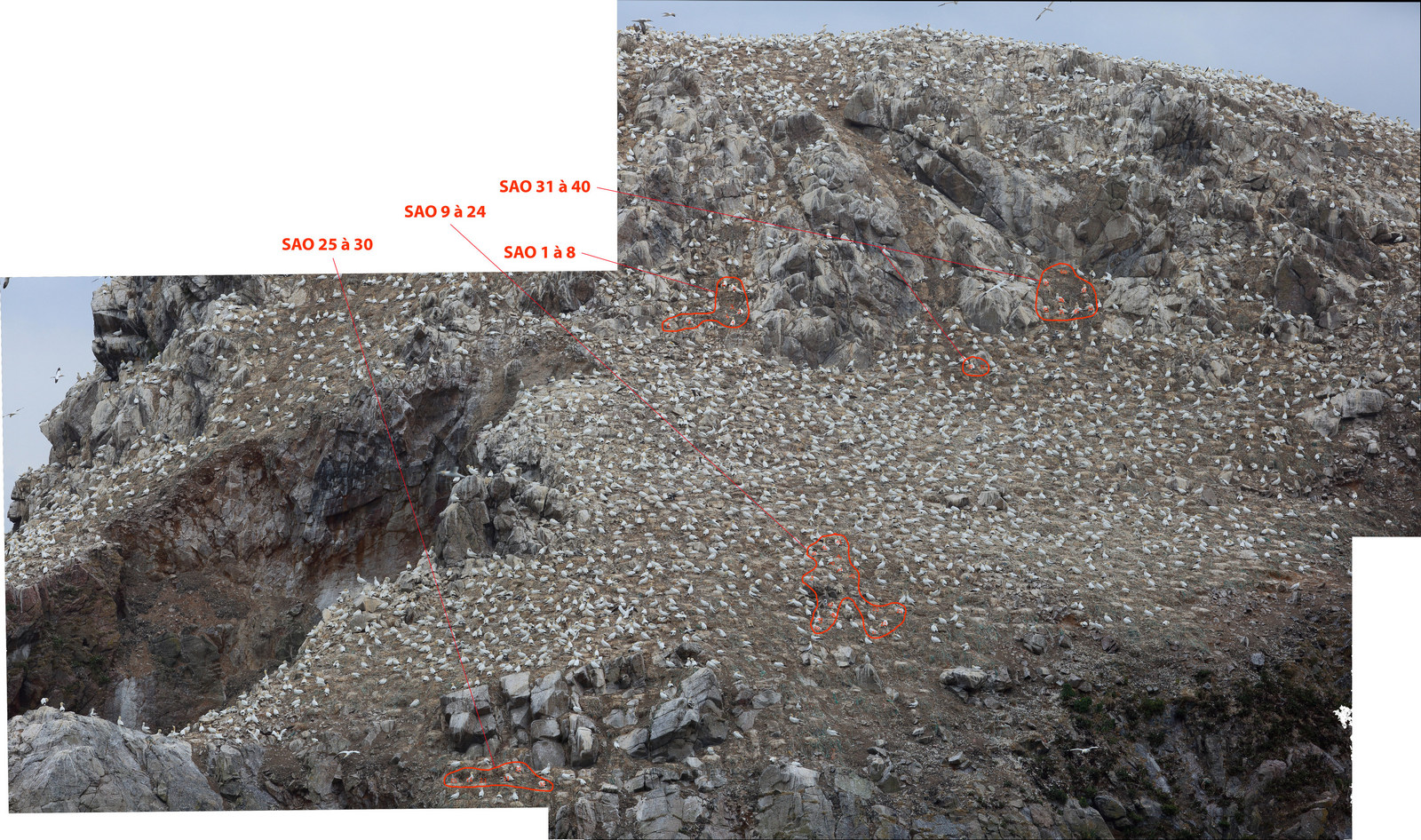

Picture S3.3. Northwest area with the 40 AOS followed indicated in red.

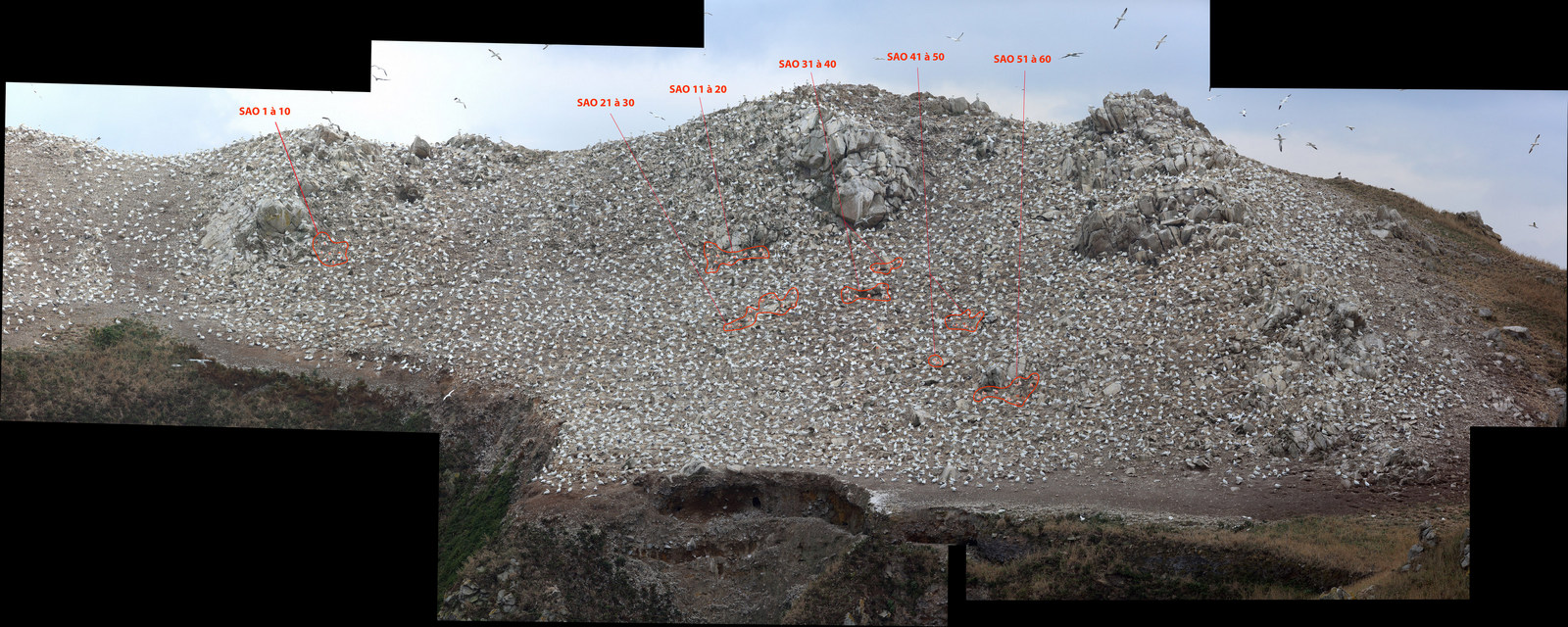

Picture S3.4. West area with the 60 AOS followed indicated in red.

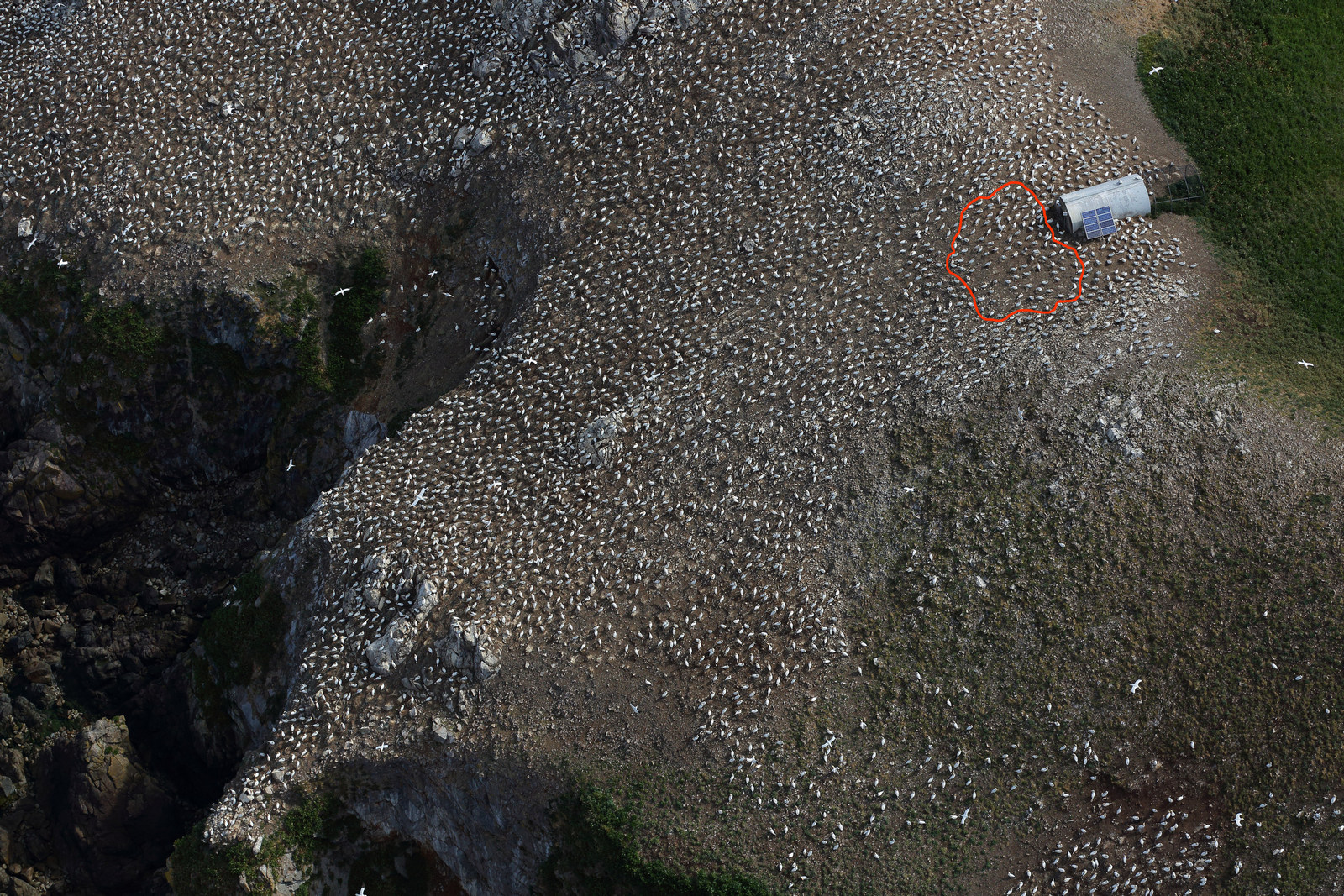

Picture S3.5. Plateau area with the 102 AOS followed. The red delineation indicated the area covered by remote CCTV.
