## SupplementaryMaterial4 for "Strong breeding colony fidelity in northern gannets following High Pathogenicity Avian Influenza Virus (HPAIV) outbreak"

Supplementary material 4: **Movement ecology of Northern gannets from the three study colonies during early chick-rearing in 2019**

Information on the movement ecology of adult breeding Northern gannets was collected and analysed using the same methodology as for the 2022 data presented in the main document.

***Table S4***. *Sample size, foraging trip characteristics, percentage of activities and colony attendance of Northern gannets nesting on Rouzic, Grassholm and Bass Rock in 2019. Results are shown as mean ± SE*

| | Bass Rock | Grassholm | Rouzic |
|---|---|---|---|
| **Total nb of equipped individuals** | 22 | 7 | 14 |
| **Nb of individuals retained** | 22 | 7 | 14 |
| **Total nb of foraging trips** | 66 | 11 | 27 |
| **Average nb of foraging trips per individual** | 3.0 ± 0.3 | 1.6 ± 0.3 | 2.0 ± 0.3 |
| **Total Nb of periods on the colony** | 35 | NA | 41 |
| **Average nb of periods on the colony per individual** | 2.7 ± 0.2 | NA | 2.4 ± 0.2 |
| **Tracking period** | 23/06/2019 – 30/07/2019 | 15/07/2019 – 17/07/2019 | 21/06/2019 – 28/06/2019 |
| **Maximal distance to the colony (km)** | 186 ± 25 | 147 ± 18 | 127 ± 12 |
| **Total distance travelled per trip (km)** | 450 ± 60 | 450 ± 78 | 328 ± 35 |
| **Trip duration (h)** | 27 ± 3 | 34 ± 8 | 20 ± 2 |
| **Time at the colony between at-sea trips (h)** | 26 ± 2 | NA | 22 ± 2 |

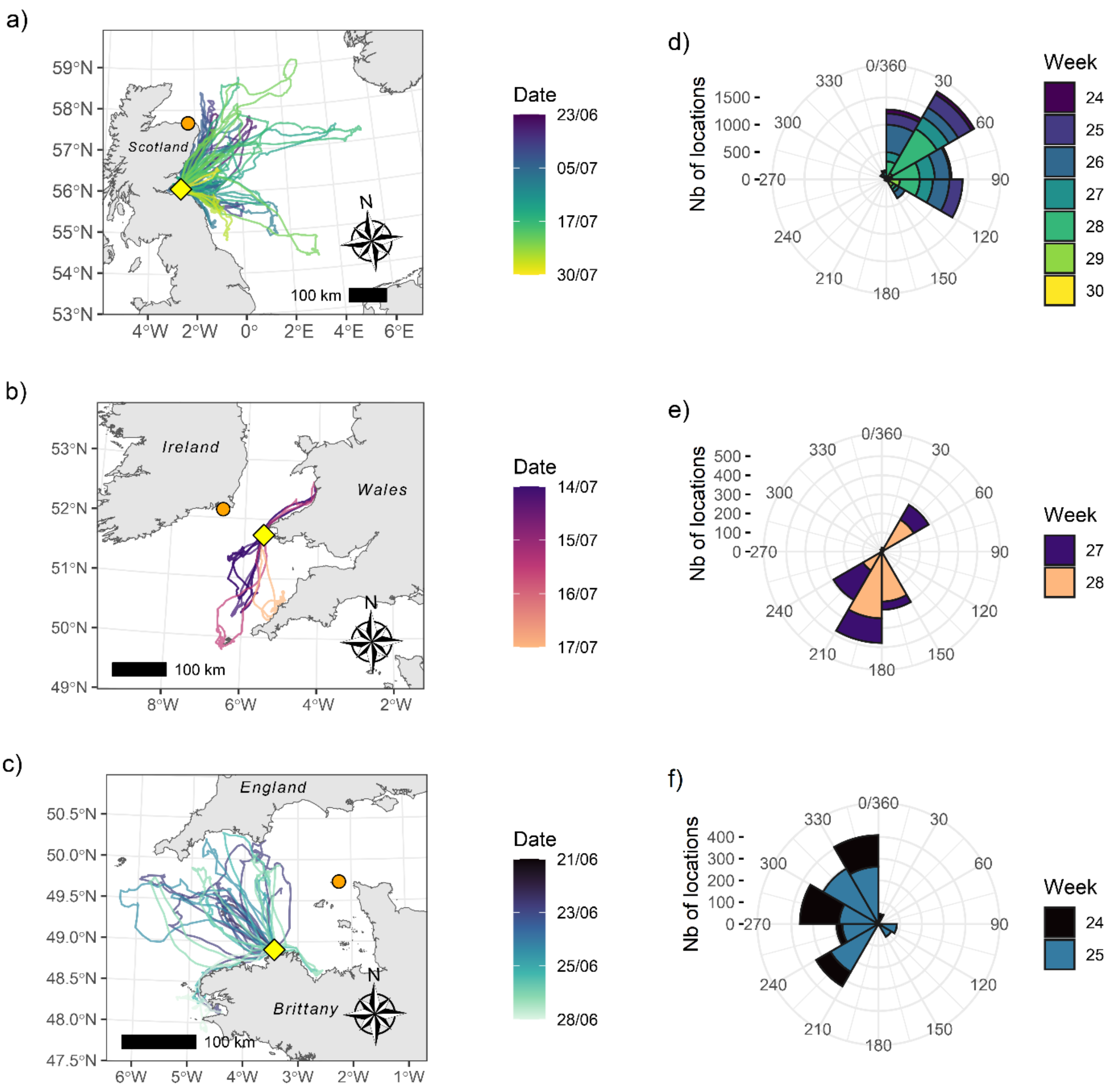


***Figure S4****. Tracks of Northern gannets nesting in 2019 on (a) Bass Rock, Scotland, (b) Grassholm, Wales and (c) Rouzic, Brittany, (yellow diamonds). Orange circles represent the closest neighbouring gannet colony from the studied colonies. Gradual changes in colour represent time in days since start of GPS tracking. (d-f) Rose diagrams showing the direction of at-sea locations. The centre of each rose diagram represents the colony location and each wedge represents the number of locations recorded in that direction over time, each colour representing one week.*
